# Transcriptional burst initiation and polymerase pause release are key control points of transcriptional regulation

**DOI:** 10.1101/275354

**Authors:** Caroline Bartman, Cheryl A. Keller, Belinda Giardine, Ross C. Hardison, Gerd A. Blobel, Arjun Raj

## Abstract

Transcriptional regulation occurs via changes to the rates of various biochemical processes. Sequencing-based approaches that average together many cells have suggested that polymerase binding and polymerase release from promoter-proximal pausing are two key regulated steps in the transcriptional process. However, single cell studies have revealed that transcription occurs in short, discontinuous bursts, suggesting that transcriptional burst initiation and termination might also be regulated steps. Here, we develop and apply a quantitative framework to connect changes in both Pol II ChIP-seq and single cell transcriptional measurements to changes in the rates of specific steps of transcription. Using a number of global and targeted transcriptional regulatory perturbations, we show that burst initiation rate is indeed a key regulated step, demonstrating that transcriptional activity can be frequency modulated. Polymerase pause release is a second key regulated step, but the rate of polymerase binding is not changed by any of the biological perturbations we examined. Our results establish an important role for transcriptional burst regulation in the control of gene expression.

## Introduction

The transcription of DNA into RNA is a highly regulated process, and many cellular decisions and responses manifest as changes in the rate of transcription. The rates of a number of transcriptional steps could be regulated in principle, but it is not clear what the main regulated steps are, in part because different experimental methods have pointed to different regulated steps. Population-averaging biochemical and sequencing assays suggest that transcription is mainly regulated at two steps: first by the recruitment of RNA polymerase II (Pol II) to the promoter of a gene, facilitated by transcription factors at enhancers and promoters, and second by polymerase pause release from the promoter proximal region facilitated by the kinase complex p-TEFb <u>(Levine et al. 2014; Jonkers and Lis 2015; Hager et al. 2009; Juven-Gershon et al. 2008; Goodrich and Tjian 2010; Core et al. 2008; Core and Lis 2008; Guenther et al. 2007; Muse et al. 2007; Henriques et al. 2013)</u>. After these two steps occur, productive elongation by Pol II takes place, generating one nascent RNA. Thus, the consensus in the field is that transcriptional regulation could occur by changing either the rate of polymerase recruitment, the rate of polymerase pause release, or both.

Single cell studies have revealed, however, that genes fluctuate between transcriptionally active and inactive states, and that the rates of these fluctuations, which we will call burst initiation and burst termination, may also be subject to regulation <u>(Bartman et al. 2016; Senecal et al. 2014; Kalo et al. 2015; Raj et al. 2006; Bahar Halpern et al. 2015)</u>. During each transcriptionally-active period or 'transcriptional burst', multiple rounds of polymerase binding and polymerase pause release can occur, while no RNAs are produced during an inactive period <u>(Raj et al. 2006; Zenklusen et al. 2008; Suter et al. 2011; Chubb et al. 2006; Bahar Halpern et al. 2015; Lionnet et al. 2011; Coleman et al. 2015; Golding et al. 2005; Dar et al. 2012)</u>. The transitions between the transcriptionally-active and -inactive states are typically slower than those of polymerase binding and polymerase pause release (hours vs. tens of minutes) <u>(Henriques et al. 2013; Cisse et al. 2013; Cho et al. 2016; Coulon et al. 2013; Jonkers et al. 2014; Tantale et al. 2016)</u>. Therefore, changes to burst initiation and burst termination may form a layer of regulation that is superimposed on top of polymerase binding and polymerase pause release.

Together, the results of population-based polymerase occupancy assays along with single cell transcriptional measurements suggest the following decomposition. Both the overall rate of polymerase recruitment and polymerase pause release are subject to biological regulation. However, overall polymerase recruitment does not consist only of binding of the polymerase to the promoter. Rather, we hypothesize that it is itself a multi-step process, consisting of first burst initiation, then polymerase binding to the promoter, then burst termination. Thus, biological regulation of the overall polymerase recruitment rate could consist of changes to the burst frequency (i.e., burst initiation or termination; frequency modulation) or burst size (i.e., polymerase binding rate; amplitude modulation). Our goal here was to discriminate these two possibilities.

Methodologically, it has proven difficult to distinguish these possibilities in part because single cell and population averaging measures of transcription give different types of information. Nascent transcript RNA FISH uses probes specific to introns of a gene of interest to measure the active-transcribing fraction (the fraction of copies of a gene making nascent transcripts), and the nascent RNA intensity (the mean fluorescence intensity of those nascent transcription sites, which is correlated to the number of transcripts recently synthesized) (Femino et al. 1998; Raj et al. 2006; Fremeau et al. 1986). (Previous publications have referred to the active-transcribing fraction as the 'burst fraction', and the nascent RNA intensity as 'burst size', but such experimental measures are in principle independent of whether or not a gene transcribes in a burst-like manner.)

In contrast, Pol II ChIP-seq, which measures average genome-wide polymerase occupancy, and NET-seq and GRO-seq, which measure average instantaneous transcriptional activity <u>(Churchman and Weissman 2011; Core et al. 2008; Kwak et al. 2013)</u>, cannot measure rates of bursting, since these methods average signal across large populations of cells. However these methods *are* able to measure genome-wide changes in polymerase pause release rate, which nascent transcript RNA FISH cannot. Changes in polymerase pause release rate can be measured by the 'Pol II traveling ratio' or 'pausing ratio', which quantify the relative amount of Pol II binding at the gene body versus near the promoter <u>(Winter et al. 2017; Rahl et al. 2010; Henriques et al. 2013)</u>. Given the complementary information provided by Pol II ChIP-seq and nascent transcript RNA FISH, combining data from these methods would allow us to directly compare changes in burst initiation and termination rates with changes in polymerase binding and polymerase pause release rates, enabling us to identify which of these steps are regulated by different perturbations.

Thus, to determine which steps of transcription are most commonly changed by biological perturbation, we developed a quantitative framework to predict how nascent transcript RNA FISH and Pol II ChIP-seq measurements would change if rates of individual steps of transcription were changed. We then performed both nascent transcript RNA FISH and Pol II ChIP-seq in the presence of a number of global and gene-specific perturbations, and compared the results to our model's predictions. We found that the alteration of two of the steps of transcription account for the majority of the quantitative changes in gene regulation. First, we found that burst initiation was the primary way in which overall polymerase recruitment to the promoter was regulated, whereas polymerase binding rate was generally not regulated by biological perturbations. Increased enhancer-promoter contact, as well as acute increases in transcription factor levels, specifically increased only the burst initiation rate, suggesting potential biochemical underpinnings of burst initiation. We also found that polymerase pause release rate is the other major regulated step of transcription. Our study supports a model of transcription in which overall polymerase recruitment is predominantly frequency modulated, by altering burst initiation rate, rather than amplitude modulated by altering polymerase binding rate; and it implicates transcriptional burst initiation as a critical control point in transcriptional regulation.

## Results

### Burst initiation is a regulated step of transcription

We set out to identify the key regulated steps of transcription by using a quantitative framework to predict how changing rates of different transcriptional steps should affect experimental measurements. First we asked whether burst initiation and termination might be regulatory steps of transcription, or whether only polymerase binding and polymerase pause release are regulated (Figure 1A). We generated two models, one including only polymerase binding and polymerase pause release as regulated steps (binding-release model, Figure 1A), and one that also incorporated burst initiation and termination steps (burst-binding-release model). (For definitions of terms used below, see Supplementary Figure 1). These models predicted that nascent transcript RNA FISH could distinguish changes in burst initiation rate from changes in rates of other transcriptional steps (Figure 1B-C, Supplementary Figure 2). Nascent transcript RNA FISH uses fluorescent probes specific to introns to quantify recent transcription in single cells (Figure 1E, left, Supplementary Figure 1). If burst initiation rate were increased, the frequency of transcriptionally-active periods would increase, resulting in an increased active-transcribing fraction of cells (Figure 1D, right). However, each active period should result in approximately the same number of RNAs produced each time, so the nascent RNA intensity would be unchanged. In contrast, a change in the polymerase binding rate was predicted to increase both the active-transcribing fraction and nascent RNA intensity (Figure 1D, left).

**Figure 1.**
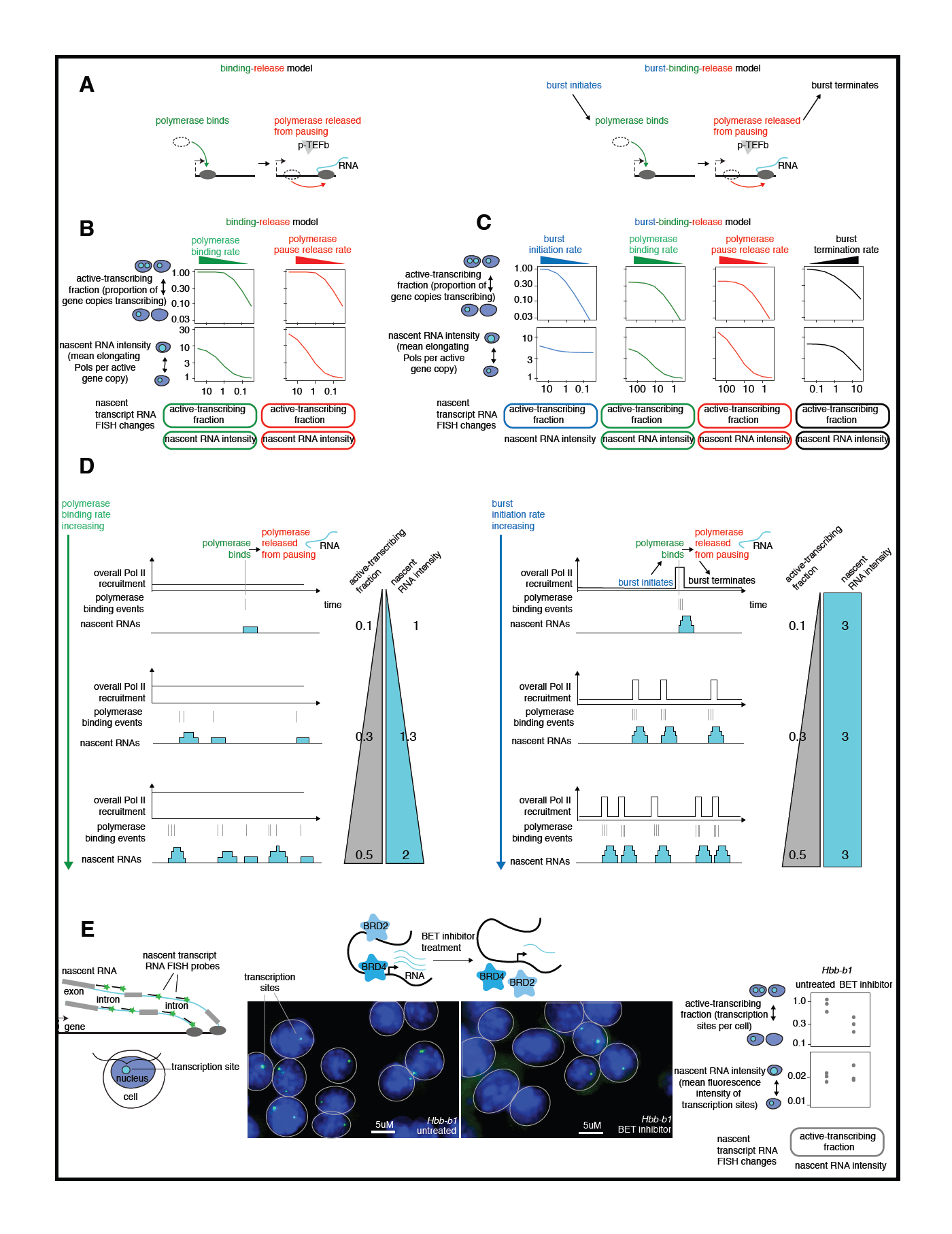
Burst initiation is a regulated step of transcription. A. Schematic of two possible models of transcription, the binding-release model and the burst-binding-release model. B. Predictions of the binding-release model for how changes in specific transcriptional rates should change nascent transcript RNA FISH measurements. C. Predictions of the burst-binding-release model for how changes in specific transcriptional rates should change nascent transcript RNA FISH measurements. D. Schematic of how changing the rate of polymerase binding is predicted to change both active-transcribing fraction and nascent RNA intensity, while changing the rate of burst initiation changes only active-transcribing fraction, while nascent RNA intensity remains unchanged. E. Left, Schematic of nascent transcript RNA FISH. Middle, nascent transcript RNA FISH images of *Hbb-b1* in G1E-ER4 cells differentiated 24 hours, untreated or with 250nM JQ1 for 60 minutes, adjusted to same brightness for all images. Right, quantification of active-transcribing fraction and nascent RNA intensity of *Hbb-b1* in the presence and absence of BET inhibitor. Each point is the mean of one biological replicate (n=3).

We first examined whether BET inhibitor treatment might regulate burst initiation rate. BET inhibitors block the transcriptional activator proteins BRD2, BRD3 and BRD4 from binding to chromatin, and may inhibit multiple facets of gene regulation including polymerase pause release and enhancer activity (Shi and Vakoc 2014; Belkina and Denis 2012; Stonestrom et al. 2016). Acute BET inhibitor treatment of the G1E-ER4 erythroid cell line decreased the active-transcribing fraction of the *Hbb-b1* gene, which codes for β-globin, without changing its nascent RNA intensity (Figure 1E). We concluded that BET inhibition reduced the burst initiation rate of *Hbb-b1*, so we can reject the simplest (binding-release) model of transcription in favor of the burst-binding-release model. This finding suggests that burst initiation can be a regulated step of transcription and thus transcriptional activity can be frequency modulated.

### The burst-binding-release model predicts experimental outcomes of changing rates of each transcriptional step

Since our previous data supported the inclusion of a burst initiation step in our transcriptional model, for the rest of this study we will examine and test the predictions of the burst-binding-release model (Figure 1B). To provide more detail on how this model connected changes in transcriptional steps to experimental predictions, a gene copy begins in a transcriptionally-unavailable state (Figure 2A). It can then transition to a transcriptionally-available state: this transition occurs faster on average if the burst initiation rate is higher, slower if the burst initiation rate is lower. If a gene is in the transcriptionally-available state, one polymerase can bind to the gene promoter, and then that polymerase can be released from pausing to produce one RNA, with the rate of each of these events determined by their respective rate constants. From either the polymerase-bound or –unbound state, the gene copy can transition back to the transcriptionally-unavailable state, with a rate defined as the rate of burst termination.

**Figure 2.**
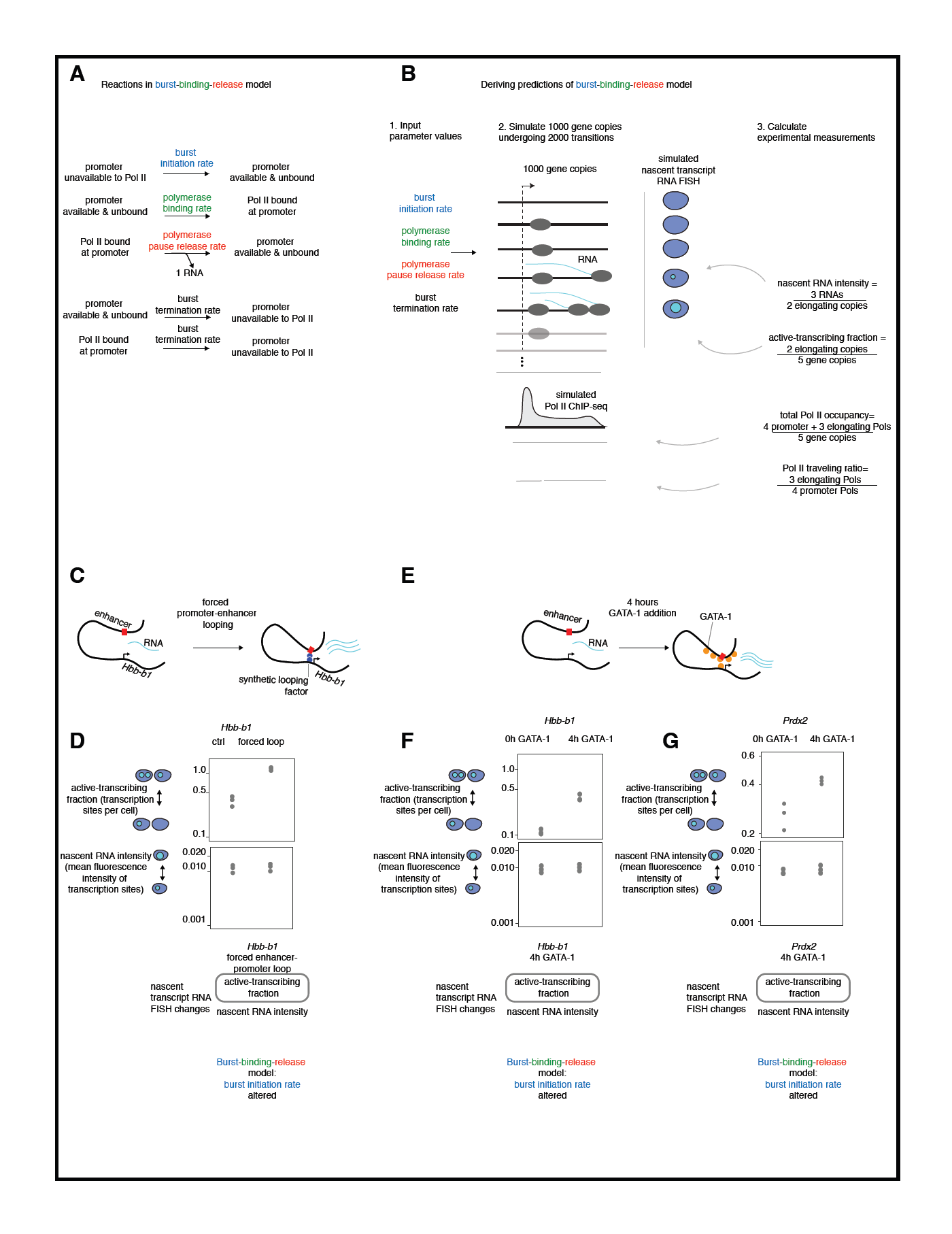
Increased enhancer-promoter looping and increased transcription factor levels regulate the rate of burst initiation. A. Equations of the burst-binding-release model of transcription. B. Schematic of how stochastic simulations of the burst-binding-release model predict the experimental outcome of changing individual transcriptional rates. C. Schematic of forced *Hbb-b1* enhancer-promoter looping. D. Top: active-transcribing fraction of *Hbb-b1* measured by nascent transcript RNA FISH; bottom: nascent RNA intensity of *Hbb-b1* measured by nascent transcript RNA FISH in G1E-ER4 cells differentiated for 9 hours, with or without expression of the looping factor. Each point is the mean of one biological replicate (n=3), data is from (Bartman et al. 2016). E. Schematic of 4 hour GATA-1 addition. F. Top: active-transcribing fraction of *Hbb-b1*; bottom: nascent RNA intensity of *Hbb-b1.* Both were measured by nascent transcript RNA FISH in G1E-ER4 cells untreated or treated with estradiol to stabilize GATA-1 for 4 hours. Each point is the mean of one biological replicate (n=3). G. Top: active-transcribing fraction of *Prdx2*; bottom: nascent RNA intensity of *Prdx2.* Both were measured by nascent transcript RNA FISH in G1E-ER4 cells untreated or treated with estradiol to stabilize GATA-1 for 4 hours. Each point is the mean of one biological replicate (n=3).

To make experimental predictions for the effects of changing each transcriptional rate, we performed stochastic simulations using a set of four values, one for each transcriptional rate in this model (Figure 2B). We simulated one thousand gene copies that over time could undergo burst initiation, polymerase binding, polymerase pause release, and burst termination. Each simulated gene copy underwent 2000 changes in state, and we recorded the state of each gene copy at 1500 regularly-spaced time intervals throughout the simulation. Every gene copy started in the transcriptionally-unavailable state; the simulation equilibrated away from the initial condition for every property, and typically converged after 100-200 time steps. The value at which it converged was used for the following analyses.

To predict Pol II ChIP-seq and nascent RNA FISH experimental outcomes from this simulated set of 1000 gene copies, we considered that each elongating polymerase produced one RNA and elongated along the gene body for a short, fixed amount of time, starting from the time in the simulation that gene underwent polymerase pause release. We then used the information of when each gene copy had promoter-bound or elongating polymerase to calculate predictions of Pol II ChIP-seq and nascent RNA FISH experimental measurements. These included active-transcribing fraction (proportion of gene copies with at least one polymerase elongating in the gene body at a given time), nascent RNA intensity (average number of elongating polymerases per transcribing gene copy), polymerase occupancy at the promoter (proportion of gene copies in the Pol II promoter-bound state), and polymerase occupancy at the gene body (average number of elongating polymerases per gene copy). Finally, we calculated Pol II traveling ratio as the ratio of mean gene body polymerase occupancy to mean promoter polymerase occupancy, and calculated total Pol II occupancy as the sum of promoter polymerase occupancy and gene body polymerase occupancy (see Supplementary Figure 1 for a glossary of experimental measurements and transcriptional steps used in this study).

We then varied the value for each of the four rates through a 1000-fold range of values, holding the other parameters constant, and performed simulations of 1000 gene copies for each set of values. With the results of all of the simulations, we could predict how changing the rate of a given transcriptional step would be predicted to change Pol II ChIP-seq and nascent transcript RNA FISH measurements.

### Enhancer-promoter looping and acute increase of transcription factor level regulate the rate of burst initiation

Given that BET inhibition reduced burst initiation rate for *Hbb-b1*, we wondered whether more gene-specific regulatory perturbations like manipulating enhancer elements or transcription factor levels also act by changing the rate of burst initiation. Contact between an enhancer and its target promoter is thought to be a critical aspect of transcriptional regulation, so we took advantage of a system to specifically increase promoter-enhancer contact at *Hbb-b1* (Figure 2C) (Deng et al. 2012). We expressed a synthetic transcription factor that binds specifically at the *Hbb-b1* promoter, fused to an LDB1 dimerization domain which dimerizes with an LDB1 protein molecule naturally bound to the *Hbb-b1* enhancer. We found that increasing promoter-enhancer contact at *Hbb-b1* increased active-transcribing fraction without changing nascent RNA intensity (Figure 2D), consistent with a change to burst initiation rate. This suggests that promoter-enhancer interactions may regulate burst initiation rate, suggesting a biochemical basis for the regulation of burst initiation.

Since transcription factor binding can facilitate enhancer-promoter looping, we sought to extend the above finding by acutely increasing the level of the key erythroid transcription factor GATA-1. GATA-1 is known to directly increase enhancer-promoter interactions for *Hbb-b1*, thus increasing transcription (Vakoc et al. 2005; Jing et al. 2008). We acutely increased levels of GATA-1 for 4 hours using a GATA-1-estradiol-receptor expressing erythroid cell line (G1E-ER4) (Figure 2E), in order to modulate transcription but minimize the indirect effects that would result from longer-term increase in GATA-1 levels (see Figure 6 for analysis of longer term treatment with more complex effects). We found that *Hbb-b1* active-transcribing fraction increased but that nascent RNA intensity was unchanged by four hours of increased GATA-1 levels (Figure 2F), demonstrating a change in burst initiation rate. An acute increase in GATA-1 levels also increased active-transcribing fraction but not nascent RNA intensity for the *Prdx2* gene (Figure 2G). Therefore GATA-1 binding affects multiple early-induced genes by changing burst initiation rate, implicating regulation of burst initiation as a broadly important mode of transcriptional regulation, and identifying increased transcription factor levels as a second biochemical correlate of burst initiation.

### Nascent transcript RNA FISH can distinguish changes in burst initiation rate from changes in polymerase binding rate

We next sought to confirm that nascent transcript RNA FISH could distinguish a change in burst initiation rate from a change in polymerase binding rate as predicted by the burst-binding-release model. We took advantage of the previously characterized drug triptolide, which specifically reduces the ability of Pol II to bind to the promoter (Figure 3A) (Jonkers, Kwak, and Lis 2014; Henriques et al. 2013; Titov et al. 2011; Vispé et al. 2009). We found that 60 minute triptolide treatment indeed reduced both active-transcribing fraction and nascent RNA intensity for all five genes examined (Figure 3B-C). These measurements conformed to the predictions of the burst-binding-release model, and were distinct from the changes observed when burst initiation rate alone was changed (Figure 3D, Figures 1 and 2). These results demonstrate that nascent transcript RNA FISH can indeed distinguish regulation of burst initiation rate from polymerase binding rate.

**Figure 3.**
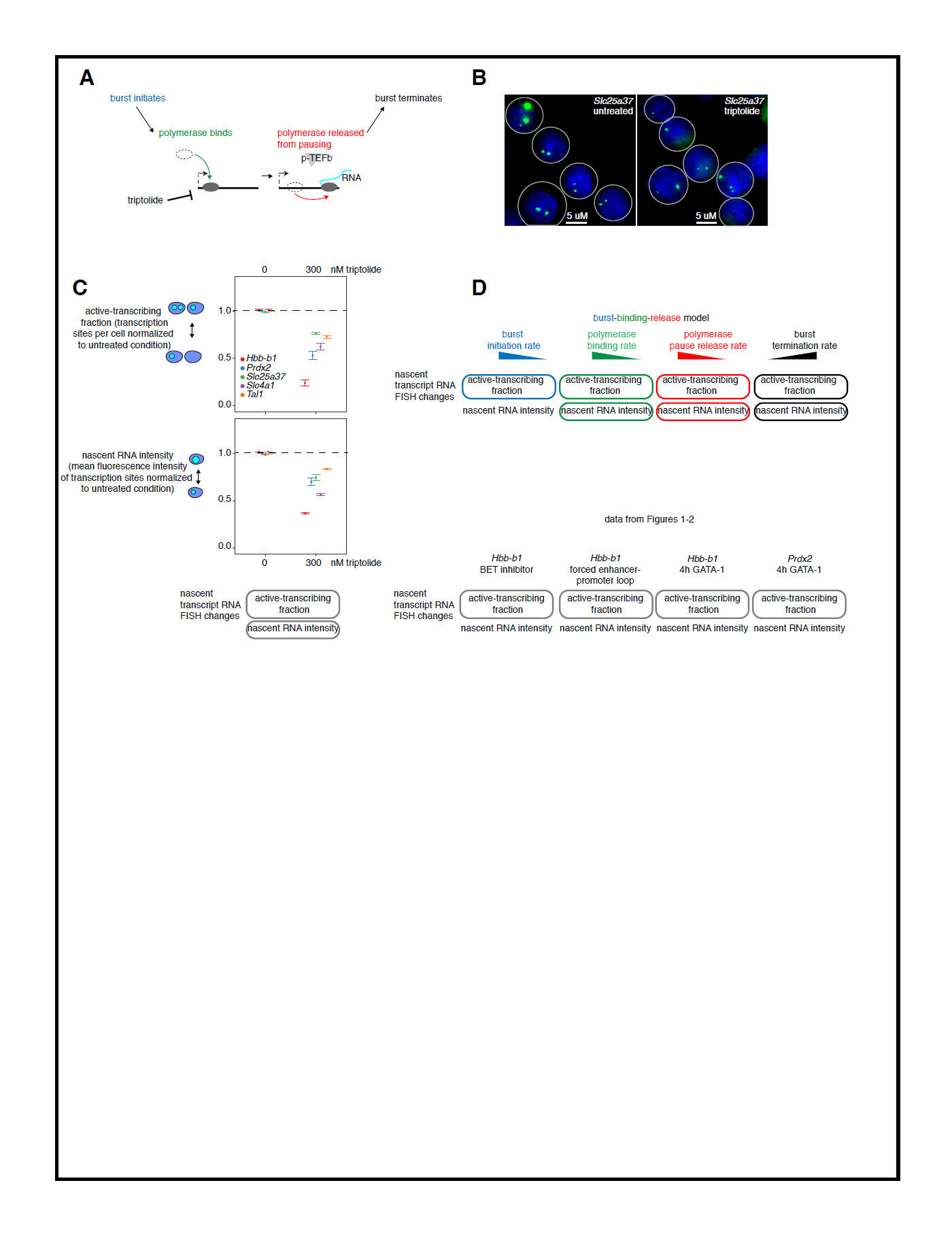
Nascent transcript RNA FISH can distinguish changes in burst initiation rate from changes in polymerase binding rate. A. Schematic of triptolide treatment. B. Nascent transcript RNA FISH images of *Slc25a37* in G1E-ER4 cells differentiated 24 hours, untreated or treated with 300nM triptolide for 60 minutes, adjusted to the same brightness. C. Active-transcribing fraction and nascent RNA intensity of 5 genes in G1E-ER4 cells differentiated for 24 hours, untreated or treated with 300nM triptolide for 60 minutes (mean of n=3 biological replicates per gene, error bars are SEM). D. Top, predictions of burst-binding-release model for how changing individual transcriptional rates is predicted to change nascent transcript RNA FISH measurements; bottom, summary of nascent transcript RNA FISH results from Figures 1 and 2 for reference.

### Pol II ChIP-seq can distinguish changes in polymerase pause release rate from changes in polymerase binding rate or burst initiation rate

We have shown that we can use the burst-binding-release model to discriminate whether changes in regulation affect the rates of either burst initiation or polymerase binding. We next wanted to use the model to detect changes to the rate of polymerase pause release, which has already been identified as a key regulatory control point in the transcriptional process. The model predicted that nascent transcript RNA FISH cannot distinguish changes in polymerase pause release rate from changes in either polymerase binding rate or burst termination rate (see Figure 1C) because all of these affect both active transcribing fraction and nascent RNA intensity.However, Pol II traveling ratio, as measured by Pol II ChIP-seq, has been used to measure polymerase pause release rate. Importantly, our model showed that changes to other rates would not affect Pol II traveling ratio at all, thus indicating that Pol II traveling ratio can be used to specifically detect changes to the rate of polymerase pause release (Figure 4A-B, Supplementary Figure 2).

**Figure 4.**
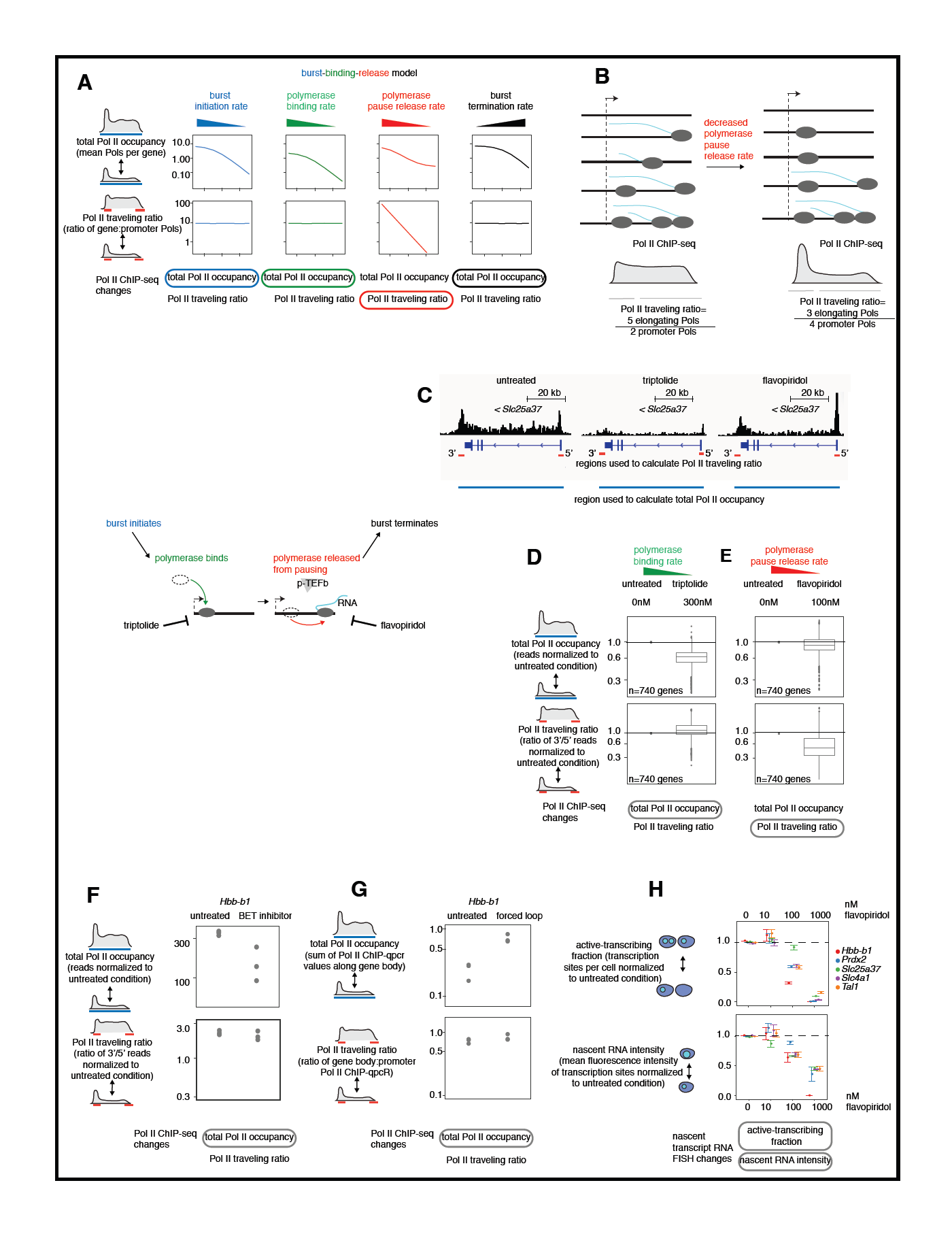
Pol II ChIP-seq can distinguish changes in polymerase pause release rate from changes in polymerase binding rate or burst initiation rate. A. Predictions of burst-binding-release model for how changing transcriptional rates is predicted to change Pol II ChIP-seq measurements. B. Schematic of how changing the polymerase pause release rate changes the Pol II traveling ratio. C. Pol II ChIP-seq tracks of *Slc25a37* in G1E-ER4 cells differentiated 24 hours, untreated or treated with 300nM triptolide or 100nM flavopiridol for 60 minutes. D. Total Pol II occupancy and Pol II traveling ratio in G1E-ER4 cells differentiated for 24 hours, untreated or treated with 60 minutes 300nM triptolide, each gene normalized to that gene in untreated cells, means of n=3 replicates each. Pol II ChIP-seq measures consider the 740 genes still transcribed after drug treatment. E. Total Pol II occupancy and Pol II traveling ratio in G1E-ER4 cells differentiated for 24 hours, untreated or treated with 60 minutes 100nM flavopiridol, each gene normalized to that gene in untreated cells, means of n=3 replicates each. Pol II ChIP-seq measures consider the 740 genes still transcribed after drug treatment. F. Top: total Pol II occupancy at *Hbb-bs* measured by Pol II ChIP-seq (from transcription start site to 1500bp after transcription end site); bottom, Pol II traveling ratio of *Hbb-bs* measured by Pol II ChIP-seq (transcription end site to 1500bp after transcription end site, divided by 750bp before to 750bp after transcription start site), in G1E-ER4 cells differentiated for 24 hours with or without 60 minutes 250nM BET inhibitor (JQ1). Each point is the mean of one biological replicate (n=3). G. Top: total Pol II occupancy at *Hbb-b1* measured by Pol II ChIP-qPCR using 4 primer pairs; bottom: Pol II traveling ratio of *Hbb-b1* measured by Pol II ChIP-qPCR using one 3' and one 5' primer pair, in G1E-ER4 cells differentiated for 9 hours with or without expression of the looping factor. Each point is the mean of one biological replicate (n=3). H. Active-transcribing fraction and nascent RNA intensity of 5 genes in G1E-ER4 cells differentiated for 24 hours, untreated or treated with the specified concentration of flavopiridol for 60 minutes (n=3 biological replicates per gene, error bars are SEM).

To verify this prediction, we treated cells with flavopiridol, which inhibits the CDK9 kinase subunit of p-TEFb, thus reducing the rate of polymerase pause release (Jonkers, Kwak, and Lis 2014; Henriques et al. 2013; Zhou, Li, and Price 2012), and compared its effects to the effects of changing the polymerase binding rate with triptolide. As predicted, reducing the elongation rate with flavopiridol strongly reduced the traveling ratio of the 740 genes still transcribed after drug treatment (Figure 4E, Supplementary Figure 3). In contrast, triptolide did not change the Pol II traveling ratio on average, though it reduced total Pol II occupancy as predicted (Figure 4C-E, Figure 4A, Supplementary Figure 3). So as predicted, Pol II ChIP-seq could distinguish a change in polymerase pause release rate from a change in polymerase binding rate.

Taking the data above together, nascent transcript RNA FISH could distinguish a change in burst initiation rate from a change in polymerase binding rate (Figure 1-3), while Pol II ChIP-seq could distinguish a change in polymerase pause release rate from a change in polymerase binding rate (Figure 4A-E). To confirm our ability to distinguish changes in individual transcriptional steps, we wanted to confirm two remaining untested predictions of the burst-binding-release model: namely, the change in Pol II ChIP-seq measurements upon a change in burst initiation rate, and the change in nascent transcript RNA FISH measurements upon a change in polymerase pause release rate. First, the model predicted that a change in burst initiation rate should result in an altered total Pol II occupancy but unchanged Pol II traveling ratio, so we performed Pol II ChIP when burst initiation rate was changed by BET inhibitor treatment or forced enhancer-promoter looping. Total Pol II occupancy was altered but Pol II traveling ratio was unchanged in both cases as the model predicted (Figure 4A, Figure 4F-G). The model also predicted that reducing the polymerase pause release rate with flavopiridol should reduce both the nascent RNA intensity and active-transcribing fraction (Figure 1C), and this prediction was also confirmed (Figure 4H).

### BET inhibition can regulate the rates of both burst initiation and polymerase pause release

So far, our observations have mostly focused on cases in which polymerase pause release was unchanged, thus allowing us to distinguish between changes to either burst initiation rate or polymerase binding rate using nascent transcript RNA FISH. However, polymerase pause release is also a major component of transcriptional regulation for many genes, and as discussed above, a change in the rate of polymerase pause release can only be distinguished using a method like Pol II ChIP-seq and not nascent transcript RNA FISH. In principle, polymerase pause release rate could change together with other rates in the model, resulting in changes to all four experimental measures, making it *a priori* hard to distinguish changes to the burst initiation rate from changes to polymerase binding rate.

For instance, we observed that BET inhibition reduced total Pol II occupancy and Pol II traveling ratio as well as active-transcribing fraction and nascent RNA intensity for *Slc25a37*, which codes for an erythroid-specific transport channel (Figure 5A-5D). Three other genes examined also displayed this behavior (Supplementary Figure 4), which was distinct from the response of *Hbb-b1* to BET inhibition. No change to a single rate alone could account for these observations.

**Figure 5.**
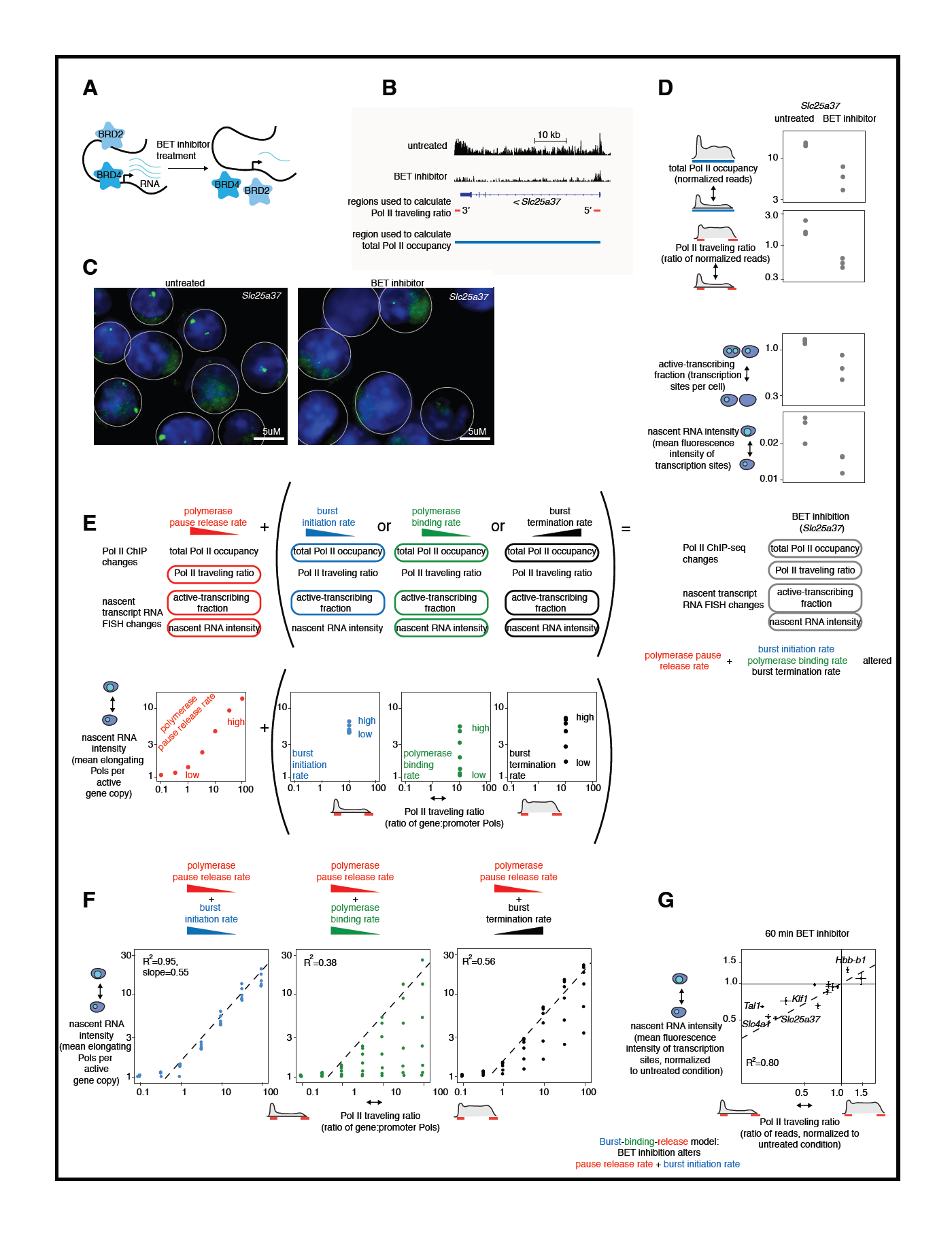
BET inhibition regulates the rates of both burst initiation and polymerase pause release. A. Schematic of BET inhibition. B. Pol II ChIP-seq tracks of Sl*c25a37* in G1E-ER4 cells differentiated for 24 hours, untreated or treated for 60 minutes with 250nM BET inhibitor (JQ1). C. Nascent transcript RNA FISH images of *Slc25a37* in G1E-ER4 cells differentiated for 24 hours untreated or treated for 60 minutes with 250nM BET inhibitor (JQ1), adjusted to same brightness for all images. D. First panel: total Pol II occupancy at *Slc25a37* measured by Pol II ChIP-seq; second panel, Pol II traveling ratio of *Slc25a37* measured by Pol II ChIP-seq; third panel: active-transcribing fraction of *Slc25a37* measured by nascent transcript RNA FISH; fourth panel: nascent RNA intensity of *Slc25a37* measured by nascent transcript RNA FISH, all in in G1E-ER4 cells untreated or treated for 60 minutes with 250nM BET inhibitor (JQ1). Each point is the mean of one biological replicate (n=3). E. Burst-binding-release model prediction for the effects of changing each transcriptional rate individually on the nascent RNA intensity and the Pol II traveling ratio. F. Burst-binding-release model prediction for the effects of changing two rates simultaneously (polymerase pause release rate and one additional rate) on the nascent RNA intensity change and the Pol II traveling ratio. The R2 value is derived from the points on the graph, each of which represents a simulation of a set of rate values in the burst-binding-release model. Each point has a different pair of values for pause release rate and the other altered rate, with the remaining two rates held constant for all points in that plot. The line log y= 0.55* log x is derived from regressing the nascent RNA intensity changes on the Pol II traveling ratio changes for the simulated points shown on the left graph (in which pause release rate + burst initiation rate change), and this line is displayed as a dotted line in all 3 graphs. G. Nascent RNA intensity vs. Pol II traveling ratio for 12 genes in G1E-ER4 cells differentiated for 24 hours treated for 60 minutes with 250nM BET inhibitor, normalized to the same gene in untreated cells. Each point is the mean of 3 biological replicates of both nascent transcript RNA FISH and Pol II ChIP-seq, error bars are SEM. Dotted line is the line log y= 0.55* log x, which is the line derived by regressing the nascent RNA intensity changes to the Pol II traveling ratio changes for the left graph in Figure 5F. R2 value for nascent RNA intensity vs Pol II traveling ratio for the measured genes is 0.80.

To begin to puzzle out which rates were changing in this more complex scenario, we first noticed that because Pol II traveling ratio was reduced, polymerase pause release rate must have been reduced because it is the only rate that can affect the traveling ratio (see Figure 4F). To match observations, however, some additional change to another rate in the model must also be occurring (Figure 5D-E). However, as shown in Figure 5E, a simultaneous change in any of either burst initiation rate or polymerase binding rate or burst termination rate would result in qualitatively similar changes to all four experimental measurements as observed, thus making it impossible to infer for any one particular gene which additional rate parameter was changing.

We thus turned to a statistical analysis aggregated over several genes to see if quantitative analysis of the relationship between different measurements could reveal which additional rate change was most likely to be occurring. Our approach was to simulate changes to both the polymerase pause release rate and an additional rate (polymerase binding rate, burst initiation rate, or burst termination rate) over a collection of values for the other rate parameters. This collection of values was intended to mimic the varying values of these rates for different genes. Strikingly, the simulations revealed that altering different additional rate parameters yielded distinguishable differences in the quantitative relationship between nascent RNA intensity and Pol II traveling ratio (Figure 5F). If we changed both burst initiation rate and pause release rate, then nascent RNA intensity and Pol II traveling ratio were predicted to display a strongly log-linear relationship, with R^2^=0.95 and a slope of approximately 0.55 (Figure 5F, left panel, each point represents a different pair of polymerase pause release rate and burst initiation rate constants, with the other two values left unchanged). This relationship, however, became far weaker if we altered polymerase binding rate together with polymerase pause release rate, with a correlation of R^2^=0.38 and many points falling below the predicted log y=0.55*log x line (Figure 5F, middle panel). Similarly, if burst termination rate is altered in addition to pause release rate, we again found a relatively low R2 value of 0.56 with many points falling below the predicted log y=0.55*log x line (Figure 5F, right panel).

We then compared the predictions to experimental measurements of response to BET inhibition across many different genes. We performed Pol II ChIP-seq and nascent transcript RNA FISH for 12 genes with and without 60 minutes of BET inhibition. We found that the 12 genes displayed a strong correlation coefficient between nascent RNA intensity change and Pol II traveling ratio change (R^2^=0.80, Figure 5G), with a log-linear relationship similar to log y= 0.55*log x. (dotted line in Figure 5G). This finding suggests that BET inhibition changes burst initiation rate and polymerase pause release rate for *Slc25a37, Slc4a1, Tal1*, and *Klf1* (Figure 5G, Supplementary Figure 4). (The remaining genes tested were either not strongly changed at all by BET inhibition, or displayed changes only in burst initiation.) It is important to note that our analysis does not definitely prove that any one gene is regulated in this manner, but rather that in aggregate, the overall pattern of regulation is most consistent with changes to burst initiation rate (in addition to polymerase pause release rate).

### Erythroid differentiation regulates the rates of both burst initiation and polymerase pause release

Since the burst-binding-release model can parse the effects of changes in multiple transcriptional rates, we next applied our approach to a more physiological perturbation, erythroid differentiation. We used the G1E-ER4 murine erythroid cell line, which expresses a GATA-1-estradiol receptor fusion. Prolonged estradiol treatment of these cells increases GATA-1 protein levels and results in both gene expression and cytological changes typical of erythroid differentiation <u>(Weiss et al. 1997)</u> (Figure 5A).

We raised GATA-1 levels by 13 hours of estradiol addition, and performed nascent transcript RNA FISH and Pol II ChIP-seq for six genes (Pol II ChIP-seq from <u>(Hsiung et al. 2016)</u>). For the three genes exhibiting significant changes (*Myc, Gata2* and *Prdx2*), we observed changes in all four measurements for each gene: total Pol II occupancy, Pol II traveling ratio, active-transcribing fraction, and nascent RNA intensity (Figure 6A-D, Supplementary Figure 5). As argued in the previous section, changes in all four of our experimental measurements could arise from a change in polymerase pause release rate plus a change in any second rate. To identify which second rate was changed by erythroid differentiation, we again examined the relationship between nascent RNA intensity change and Pol II traveling ratio change (Figure 5E). Similar to the case of BET inhibition, erythroid differentiation produced changes in nascent RNA intensity and Pol II traveling ratio which fell along the log y= 0.55* log x line (Figure 5F). Additionally, genes fell into a strong log-linear relationship with little spread, again with the correlation coefficient R^2^=0.80. These findings suggest that erythroid differentiation changes both burst initiation rate and polymerase pause release rate for *Myc, Prdx2* and *Gata2* (Figure 6F, Supplementary Figure 5).

**Figure 6.**
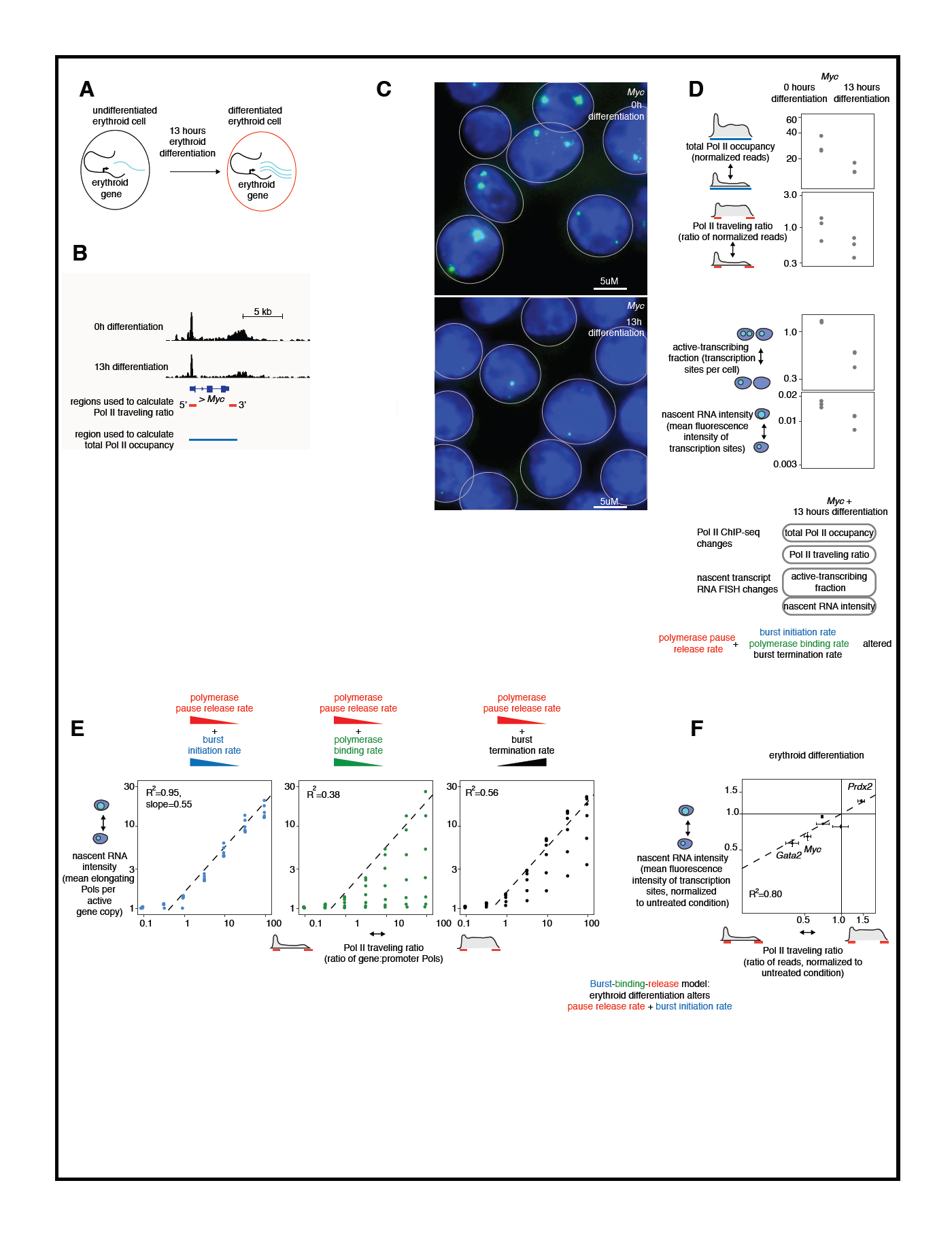
Erythroid differentiation regulates the rates of both burst initiation and polymerase pause release. A. Schematic of erythroid differentiation. B. Pol II ChIP-seq tracks of *Myc* in G1E-ER4 cells undifferentiated or differentiated for 13 hours. Data from (Hsiung et al. 2016). C. Nascent transcript RNA FISH images of *Myc* in G1E-ER4 cells undifferentiated or differentiated for 13 hours, adjusted to same brightness for all images. D. First panel: total Pol II occupancy at *Myc* measured by Pol II ChIP-seq; second panel, Pol II traveling ratio of *Myc* measured by Pol II ChIP-seq; third panel: active-transcribing fraction of *Myc* measured by nascent transcript RNA FISH; fourth panel: nascent RNA intensity of *Myc* measured by nascent transcript RNA FISH. All experiments in G1E-ER4 cells differentiated for 0 or 13 hours by addition of estradiol. Each point is the mean of one biological replicate (n=3). Pol II ChIP-seq data from (Hsiung et al. 2016). E. Burst-binding-release model prediction for the effects of changing two rates simultaneously (polymerase pause release rate and one additional rate) on the nascent RNA intensity change and the Pol II traveling ratio. The R2 value is derived from the points on the graph, each of which represents a simulation of a set of rate values in the burst-binding-release model. Each point has a different pair of values for pause release rate and the other altered rate, with the remaining two rates held constant for all points in that plot. The line log y= 0.55* log x is derived from regressing the nascent RNA intensity changes on the Pol II traveling ratio changes for the simulated points shown on the left graph (in which pause release rate + burst initiation rate change), and this line is displayed as a dotted line in all 3 graphs. F. Nascent RNA intensity vs. Pol II traveling ratio for 6 genes in G1E-ER4 cells differentiated for 13 hours, normalized to the same gene in undifferentiated cells. Each point is the mean of 3 biological replicates of both nascent transcript RNA FISH and Pol II ChIP-seq, error bars are SEM. Dotted line is the line log y= 0.55* log x, which is the line derived by regressing the nascent RNA intensity changes to the Pol II traveling ratio changes for the left graph in Figure 5F. R2 value for nascent RNA intensity vs Pol II traveling ratio for the measured genes is 0.80. Pol II ChIP-seq data from (Hsiung et al. 2016).

### Measuring genome-wide effects of BET inhibition and erythroid differentiation on burst initiation and polymerase pause release rates

Our results suggest that both BET inhibition and erythroid differentiation chiefly alter burst initiation rate and polymerase pause release rate, and seemed not to strongly affect polymerase binding rate or burst termination rate (Figures 5 and 6). If we assume that these two perturbations only alter burst initiation and polymerase pause release rates for ALL genes, we can use the genome-wide nature of Pol II ChIP-seq data to infer quantitative changes in these two rates for all genes. Under this assumption, any change in total Pol II occupancy is attributable solely to a change in burst initiation rate, and any change to Pol II traveling ratio is attributable solely to a change in polymerase pause release rate (Figure 7A).

**Figure 7.**
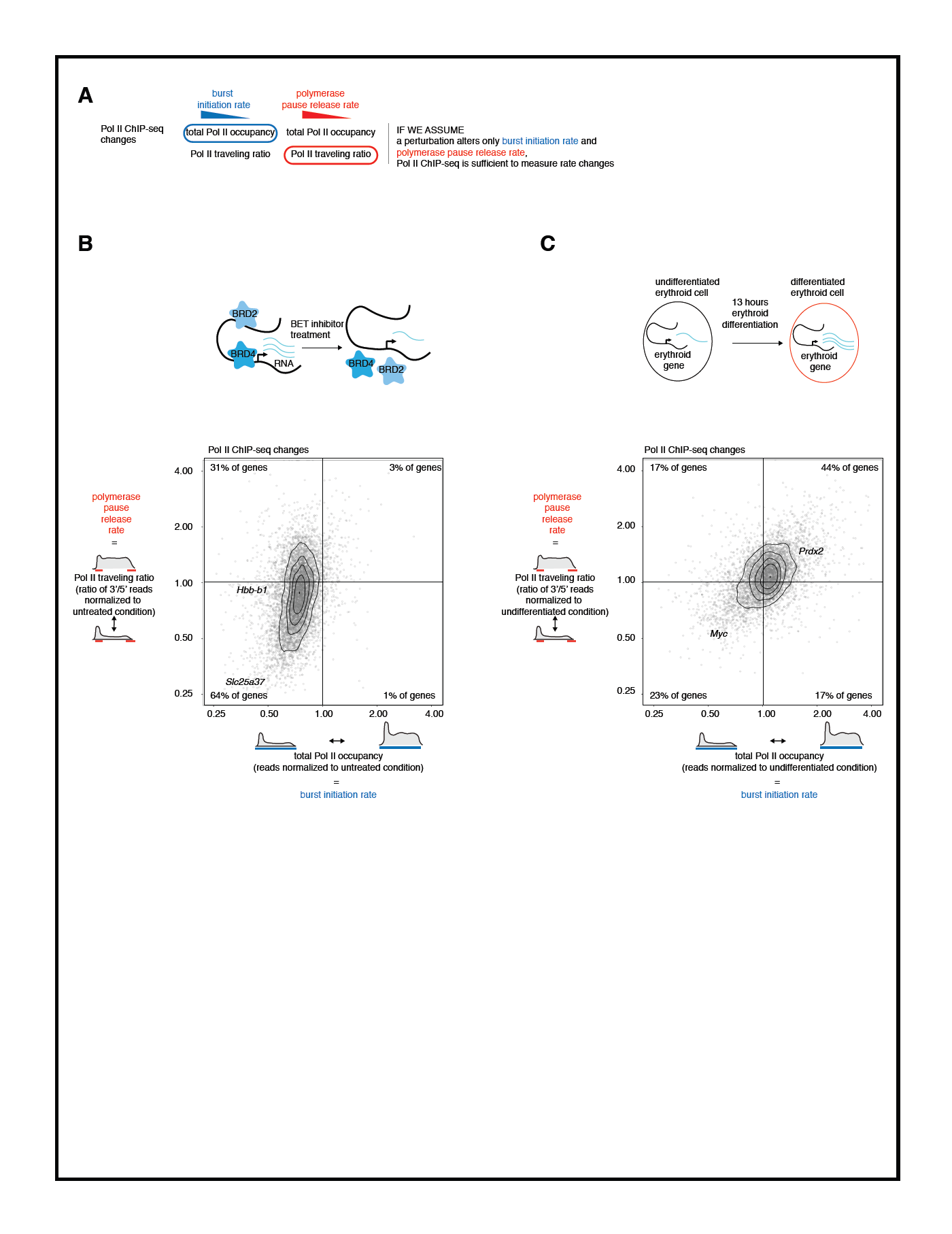
Measuring genome-wide effects of BET inhibition and erythroid differentiation on burst initiation and polymerase pause release rates. A. Predictions of burst-binding-release model: Pol II ChIP-seq can measure changes in burst initiation rate and polymerase pause release rate genome-wide if only these two rates are changed for all genes. B. Pol II traveling ratio and total Pol II occupancy for all transcribed genes (n=6609) in 24h differentiated G1E-ER4 cells with 60min 250nM JQ1, normalized to the same gene in untreated cells. Each point is the mean of n=3 biological ChIP-seq replicates (same experiments as shown in Figure 2D-E, Figure 3D-E, Figure 5D-E). C. Pol II traveling ratio and total Pol II occupancy for all transcribed genes (n=4933) in 13h differentiated G1E-ER4 cells, normalized to the same gene in undifferentiated cells. Each point is the mean of n=3 biological ChIP-seq replicates (same experiments as shown in Figure Figure 6B and 6K). Pol II ChIP-seq data from (Hsiung et al. 2016).

Assuming that BET inhibitor treatment only changes burst initiation rate and polymerase pause release rate, we found that most genes exhibited a modest but significant reduction in burst initiation rate, while many also had a reduced polymerase pause release rate (Figure 7B). Thus 31% of genes exhibit a reduction in burst initiation rate only (upper left quadrant), while 64% exhibit a reduction in both burst initiation rate and polymerase pause release rate (lower left quadrant). Many studies have suggested that the most important effects of BET inhibition (and BRD4 in particular) are on polymerase pause release (Bartholomeeusen et al. 2012; Yang et al. 2005; Jang et al. 2005; Winter et al. 2017), but our data suggests that BET proteins also play an important and almost global role in burst initiation. We also found that the genome-wide effect of BET proteins on transcription is very consistent regardless of cellular context: both differentiated and undifferentiated erythroid cells displayed a similar genome-wide response to BET inhibition, with strong changes in burst initiation rate and variable effect on polymerase pause release rate (Supplementary Figure 6).

We also examined the genome-wide changes in polymerase pause release rate and burst initiation rate produced by 13 hours of erythroid differentiation (Figure 7C). As shown in Figure 6 and Supplementary Figure 5, differentiation decreased both rates for *Myc*, while increasing both rates for *Prdx2*. Differentiation produced significant increases and decreases in both burst initiation and polymerase pause release rates genome-wide, with the most genes (44%) experiencing both increased burst initiation rate and increased polymerase pause release rate. Thus it appears that different perturbations result in different genome-wide transcriptional changes, but that these changes remained similar irrespective of context. (Note again that the conclusions of this figure can only be supported if BET inhibition and erythroid differentiation do not alter polymerase binding rate or burst termination rate.)

### Discussion

We sought to determine which steps of transcription are biologically regulated. To this end, we developed and validated a model of transcriptional regulation that maps changes in rates of polymerase binding, polymerase pause release, burst initiation and burst termination to a combination of population-based and single cell transcriptional measurements. This framework revealed that for biological perturbations we tested, overall polymerase recruitment is chiefly frequency modulated, specifically by changing the rate of burst initiation, rather than amplitude modulated by changing the rate of polymerase binding. Polymerase binding rate was not altered by the perturbations we examined except by triptolide, which is a known inhibitor of polymerase binding. Many previous studies have not separated changes in burst initiation rate from changes in polymerase binding rate because both result in less polymerase reaching a gene, leading to a finding of reduced total Pol II occupancy by population-averaging methods. We used nascent transcript RNA FISH to separate changes in these two rates and found that burst initiation is a key regulatory step, but that polymerase binding may not be. Further, Pol II ChIP-seq information combined with nascent transcript RNA FISH showed that polymerase pause release is a second key regulated step of transcription. Our findings also demonstrate the utility of combining population based and single cell transcriptional measurements.

Our results point primarily to burst initiation rate as opposed to polymerase binding rate as a key regulated step for most genes examined during the biological perturbations of our experiments. This does not mean that polymerase binding is never rate limiting. Indeed, polymerase binding rate is in fact rate limiting in the context of triptolide treatment. The finding that polymerase binding rate is not typically altered by biological perturbations corresponds well to recent single molecule imaging studies of Pol II: the authors found that even when a gene was lowly transcribed, many molecules of polymerase clustered near its promoter (Cho et al. 2016; Cisse et al. 2013). Since only one polymerase can bind a promoter at a time (Shao and Zeitlinger 2017), it is possible that polymerase binding typically occurs with a very high rate in all conditions.

It will be essential in future to better characterize the biochemical underpinnings of transcriptional bursting. An intriguing hypothesis with some experimental support is that burst initiation is related to enhancer-promoter looping. Here, we have shown that for *Hbb-b1*, increasing enhancer-promoter contact specifically increases burst initiation, consistent with other studies that have isolated looping from other potential enhancer activities <u>(Bartman et al. 2016; Fukaya et al. 2016)</u>. It is possible that each time an enhancer contacts a promoter, the gene undergoes burst initiation. This hypothesis is supported by a recent paper showing that enhancer-promoter looping occurs before every transcriptional burst (Chen, Fujioka, and Gregor 2017). Burst initiation could also be controlled by events independent from enhancer-promoter contact, such as transcription factor binding to the promoter. Mechanistic experiments will be important to understand this key regulatory step of transcription.

Our experiments with BET inhibitor treatment show that BET proteins can regulate both burst initiation rate and polymerase pause release rate. Many papers in the literature have suggested that a chief transcriptional effect of BET inhibition was to inhibit polymerase pause release, and we also found that this is a dominant effect (Figure 7) (Bartholomeeusen et al. 2012; Yang et al. 2005; Jang et al. 2005; Winter et al. 2017). However, we also observed significant effects of BET inhibitors on burst initiation rate. Indeed, BET inhibition of *Hbb-b1* only altered burst initiation rate and did not change polymerase pause release rate. Thus additional mechanisms of BET proteins such as influencing chromatin looping and enhancer activity are likely to also be globally important for modulating gene expression (Shi and Vakoc 2014; Belkina and Denis 2012; Stonestrom et al. 2016).

Strikingly, we found that BET inhibition affected different steps of transcription for different genes. It had been previously observed that some genes were totally insensitive to BET inhibition; but we also observed one gene with a reduced burst initiation rate (*Hbb-b1*) while others had reduced polymerase pause release rate and burst initiation rate (*Slc25a37, Slc4a1, Tal1, Klf1*). An important future direction will be to explore why different genes respond differently to the same perturbation.

More generally, our study demonstrates the utility of a model-based approach to identifying rate limiting steps in transcription. In particular, previous studies of the burst initiation and termination phases has been largely phenomenological, characterized primarily by observables like active-transcribing fraction and nascent RNA intensity (Senecal et al. 2014; Dar et al. 2012; Mateos-Langerak et al. 2016; Octavio, Gedeon, and Maheshri 2009). It has, however, proven difficult to discern any general rules or principles from these studies. Our study suggests that this may be due to the fact that these experimental observables can be convolved in counterintuitive ways; indeed, there is no reason *a priori* to believe that such observables map one-to-one to particular biological processes. For example, our study suggests that nascent RNA intensity was not independently regulated; rather, active-transcribing fraction and nascent RNA intensity were both altered when any one of the rates of polymerase binding, polymerase pause release, or burst termination are changed (Figures 1,3,4). By using a model-based approach informed by a combination of data types, we instead were able to interpret these observables in terms of parameters of a simple model of transcription, thus revealing a more consistent picture over a variety of perturbations.

## Experimental Procedures

### Murine cell culture, infection and sorting

G1E-ER4 cells were cultured and differentiated as described (Weiss, Yu, and Orkin 1997). All G1E-ER4 cells used were from Blobel lab. For flavopiridol experiments, the noted concentration of flavopiridol (either 10nM, 100nM, or 1uM) was added for 60 minutes. For triptolide experiments, the noted concentration of flavopiridol (either 10nM, 100nM, 300nM, or 1uM) was added for 60 minutes. For forced looping experiments, cells were infected with the MIGR-1 retrovirus expressing only GFP or mZF-SA followed by an IRES element and GFP (Deng et al. 2012). Infections were performed as described (Tripic et al. 2009). Cells were expanded for two days and sorted using a BD FacsAria to purify GFP+ infected cells from control and mZF-SA samples. Finally, estradiol was added for 9 hours and transcription was measured by FISH or ChIP-qPCR.

### Chromatin Immunoprecipitation

We performed ChIP as previously described (Letting et al. 2004), using the N-20 Pol II antibody (Santa Cruz sc899). ChIP-qPCR was performed with Power SYBR Green (Invitrogen). For ChIP-sequencing, library construction was performed using Illumina’s TruSeq ChIP sample preparation kit (Illumina, catalog no. IP-202-1012) according to manufacturer’s specifications with the addition of a size selection using SPRIselect beads (Beckman Coulter, catalog no. B23318) prior to PCR amplification. Library size was determined (average ˜340 bp) using the Agilent Bioanalyzer 2100, followed by quantitation using real-time PCR using the KAPA Library Quant Kit for Illumina (KAPA Biosystems catalog no. KK4835). Libraries were then pooled and sequenced on the Illumina NextSeq 500 using Illumina sequencing reagents according to manufacturer’s instructions.

### ChIP-sequencing analysis

All ChIP-seq data was generated for this study, except the 0 and 13h G1E-ER4 differentiation (Figure 4), which was from (Hsiung et al. 2016). We used bcl2fastq2 to convert and demultiplex the reads. We also applied fastQC to get read stats, Bowtie (0.12.8) to map reads to the mm9 genome (multiple and unique mappings), Samtools to convert SAM to BAM and get stats, MACS (1.3.7.1) to make wiggle files, and used wigToBigWig to convert wiggle files to bigWig files.

We then performed bigWigAverageOverBed to find Pol2 binding signal at gene body region, (transcription start site to 1500 base pairs downstream of transcription end site),TSS region (750 bp upstream of transcription start site to 750bp downstream), and TES region (transcription end site to 1500bp downstream of transcription end site). We then selected only the genes with strong pol2 binding in all regions in all experimental conditions, so that we should never be examining genes that aren’t transcribed (potentially resulting in dividing by 0). We did this by setting arbitrary cutoff values on the bigWigAverageOverBed results, and results were robust to changes in cutoff. We then corrected to number of aligned reads per sample. We also performed a background correction by performing bigWigAverageOverBed in three 100kb, transcription-free genomic regions, and dividing by the background signal in those regions (this changed values by a maximum of +/- 15%, but resulted in biological replicates correlating to each other more strongly). Pol II traveling ratio was computed by the ratio of TES Pol II signal to TSS Pol II signal. The only exception to this pipeline was the flavopiridol and triptolide datasets: the global transcriptional reduction achieved by these compounds resulted in a lot less material pulled down by ChIP and thus a smaller library size, so in these cases we did not normalize to library size. (Note that the only metric influenced by this change was total Pol II occupancy; Pol II traveling ratio is effectively internally normalized in each sample since it is TES divided by TSS, so library size correction has no effect on this metric).

### Single-molecule RNA FISH imaging

We performed single-molecule RNA FISH as described previously (Raj et al. 2006; Femino et al. 1998). Briefly, we fixed cells in 1.85% formaldehyde for 10min at room temperature, and stored them in 70% ethanol at 4 degrees C until imaging. We hybridized pools of FISH probes to samples, followed by DAPI staining and wash steps performed in suspension. Samples were cytospun onto slides for imaging on a Nikon Ti-E inverted fluorescence microscope using a 100x Plan-Apo objective (numerical aperture of 1.43), a cooled CCD camera (Pixis 1024B from Princeton Instruments), and filter sets SP102v1 (Chroma), SP104v2 (Chroma), and 31000v2 (Chroma) for Cy3, Cy5, and DAPI, respectively. Custom filter (Omega) was used for Alexa594. We took 45 optical z-sections at intervals of 0.35 microns, spanning the vertical extent of cells, with 1s exposure time for Cy3, Cy5, and Alexa594, and 35ms for DAPI.

### Image Analysis

We manually segmented boundaries of cells from bright field images and localized RNA spots using custom software written in MATLAB (Raj et al. 2010), with subsequent analyses performed in R. Transcription sites were identified by bright nuclear intron spots. Alleles co-transcribing human g-globin and b-globin are identified by colocalization of transcription sites.

### Mathematical modeling

Mathematical models were constructed and simulations were performed in Matlab using Gillespie’s stochastic simulation algorithm (Gillespie 1976). For the burst-binding-release model, genes could be in three states (closed, open, polymerase bound). A gene in the closed (off, non-burst initiated) state transitions to the open (bursting) state at the burst initiation rate, while a gene in the open state can have polymerase bind the promoter with the polymerase binding rate. Once a polymerase is bound, that polymerase can be released to elongation with the rate of polymerase release from pausing, and the promoter thus returns to the open and unbound state. From either the open unbound or open polymerase bound state, the gene can transition to the off state with the rate of burst termination. We varied each of these rates through a 1000 fold range of values. We expect that in general, the rates of polymerase binding and polymerase release tend to be greater than the rate of burst initiation, since this must be true to observe more than one RNA produced in a burst. Thus, we selected rates of (0.03,0.1,0.3,1,3,10,30) for the burst initiation and burst termination rates, and rates of (0.3,1,3,10,30,100,300) for the polymerase binding and polymerase pause release rates. For every set of four rates, we simulated 1000 gene copies, which were allowed to proceed through 2000 changes in state. We recorded the state of each gene copy at 1500 time intervals. The simulation equilibrated away from the initial condition (every gene copy started in the ’off’ state) for every property within this time window, typically after ˜100-200 time steps, and the value at which it converged was used for the following analyses. We then allowed each elongating polymerase to produce RNA and elongate along the gene body for a short, fixed amount of time, starting from the time that gene switched from the polymerase bound to the open state. We then used the above information to calculate: active-transcribing fraction (proportion of gene copies with at least 1 polymerase elongating in the gene body at a given time); nascent RNA intensity (average number of elongating polymerases on one gene for gene copies with at least 1 polymerase elongating in the gene body at a given time), polymerase binding signal at promoter (proportion of gene copies in the pol2 bound state), polymerase binding signal at gene body (average number of Pol molecules in the gene body). We calculated Pol II traveling ratio as gene body polymerase signal divided by promoter polymerase signal, and calculated total Pol II occupancy as promoter polymerase signal plus gene body polymerase signal.

For the binding-release model, genes could be in two states, either bound with polymerase or unbound. A gene in the unbound state can have polymerase bind the promoter with the polymerase binding rate. Once a polymerase is bound, that polymerase can be released to elongation with the rate of polymerase release from pausing, and the promoter thus returns to the open and unbound state. We varied each of these rates through a 1,000 fold range of values (0.3,1,3,10,30,100,300 for each rate). For every set of four rates, we simulated 1000 gene copies, which were allowed to proceed through 2000 changes in state. We recorded the state of each gene copy at 1500 time intervals. The simulation equilibrated away from the initial condition (every gene copy started in the ’off’ state) for every property within this time window, typically after ˜100-200 time steps, and the value at which it converged was used for the following analyses. We then allowed each elongating polymerase to produce RNA and elongate along the gene body for a short, fixed amount of time, starting from the time that gene switched from the polymerase bound to the open state. We calculated polymerase occupancy, polymerase traveling ratio, active-transcribing fraction, and nascent RNA intensity as described above.

| Rate Value | Burst Initiation Rate | Polymerase Binding Rate | Polymerase Pause Release | Burst Termination |
| --- | --- | --- | --- | --- |

|  |  |  | Rate | Rate |
| --- | --- | --- | --- | --- |
| Values tested<br>(binding-release model) | NA | 0.3,1,3,10,30,10<br>0,300 | 0.3,1,3,10,30,10<br>0,300 | NA |
| Values tested<br>(burst-binding-release model) | 0.03,0.1,0.3,1,3,<br>10,30 | 0.3,1,3,10,30,10<br>0,300 | 0.3,1,3,10,30,10<br>0,300 | 0.03,0.1,0.3,1,3,<br>10,30 |

### Plotting and graphics

We used R packages dplyr and ggplot2 to produce nearly all figures, followed by cosmetic adjustments in Adobe Illustrator.

## Acknowledgements

The authors thank members of the Blobel and Raj labs for careful reading and suggestions, especially Sara Rouhanifard and Vivek Behera. This work was supported by an NSF GRFP fellowship to C.R.B., NIH New Innovator Award 1DP2OD008514 to A.R., 1RO1 HL119479 to G.A.B., NIH NIDDK R01 DK054937 to G.A.B.,NIH NIDDK R24 DK106766 to G.A.B. and R.C.H., and UO1 129998 to A.R. and G.A.B.

---

**Supplementary Figure 1 (related to Figure 1):**
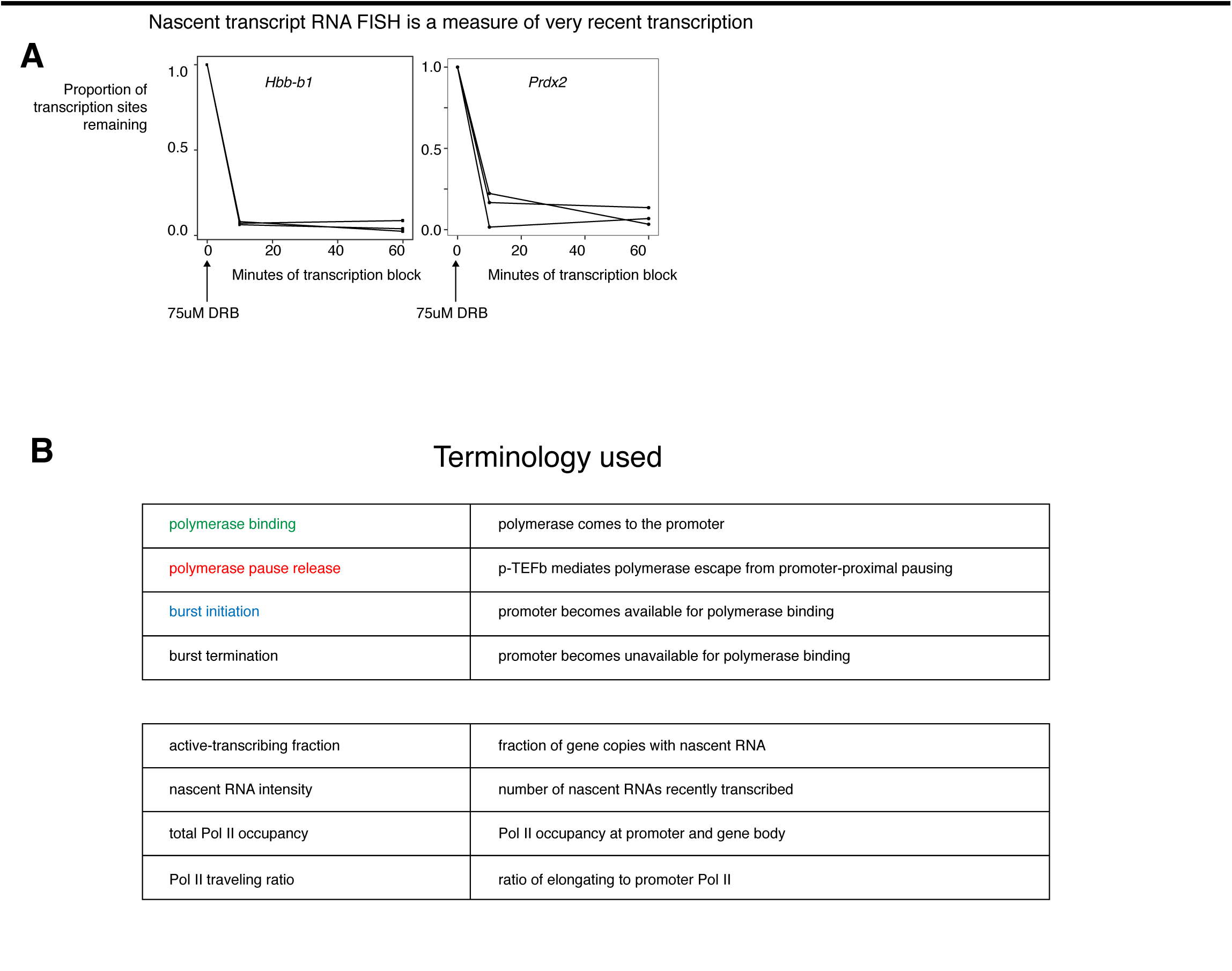
A. Active-transcribing fraction in response to 75uM DRB, measuring half-life of transcription sites (n=3 biological replicates per gene). B. Terms for transcriptional steps and experimental measures.

**Supplementary Figure 2 (related to Figure 1):**
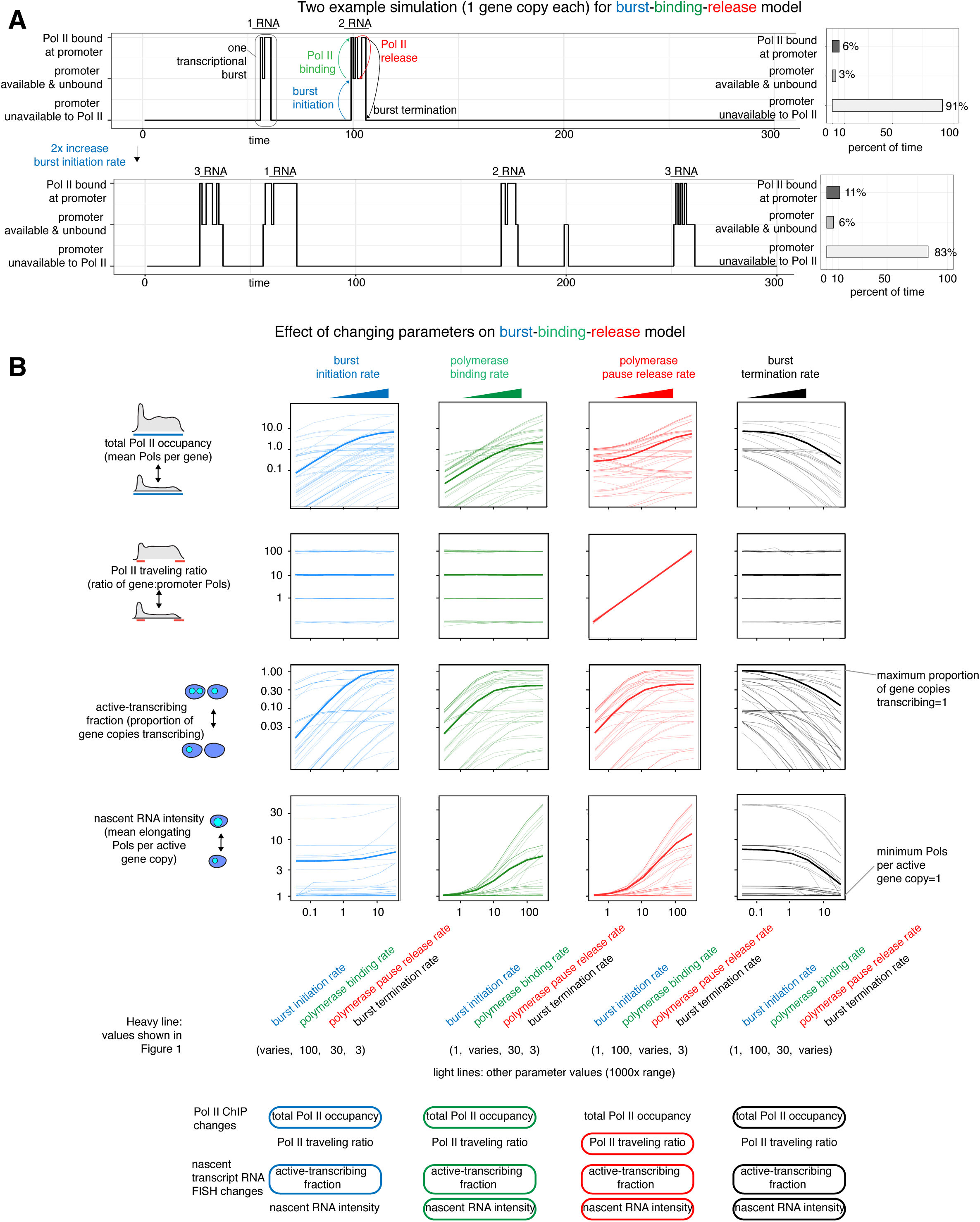
A. Example traces from simulation of one gene copy in burst-binding-release model. B. Effect of varying parameters on burst fraction, burst size, total Pol II occupancy and Pol II traveling ratio predicted by the burst-binding-release model.

**Supplementary Figure 3 (related to Figure 4):**
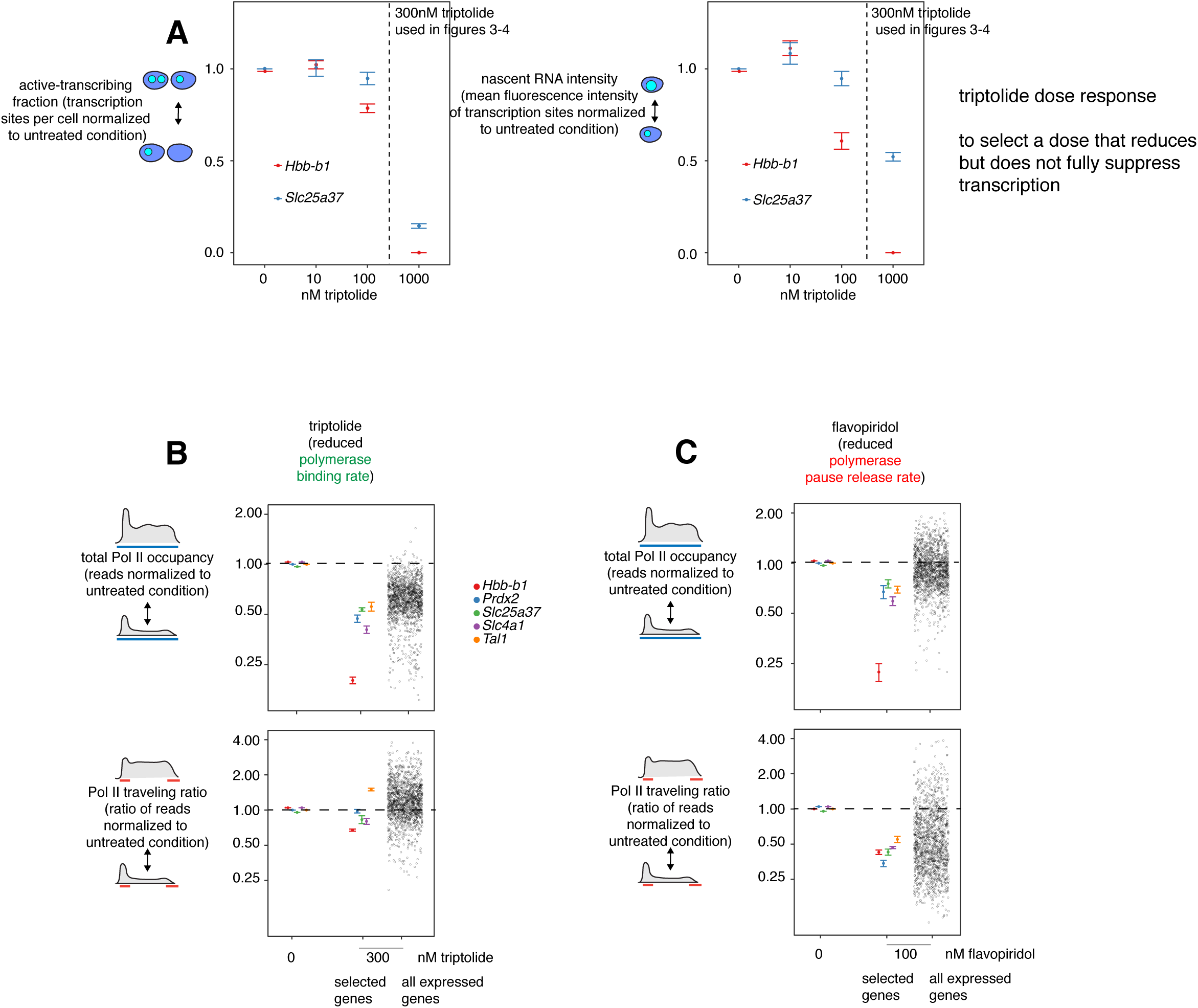
A. Active-transcribing fraction and nascent RNA intensity in response to 60min triptolide (each point= mean of 3 biological replicates per gene). B. Total Pol II occupancy change or Pol II traveling ratio change in response to 60min triptolide (each point=mean of 3 biological replicates) C. Total Pol II occupancy change or Pol II traveling ratio change in response to 60min flavopiridol (each point=mean of 3 biological replicates).

**Supplementary Figure 4(related to Figure 5):**
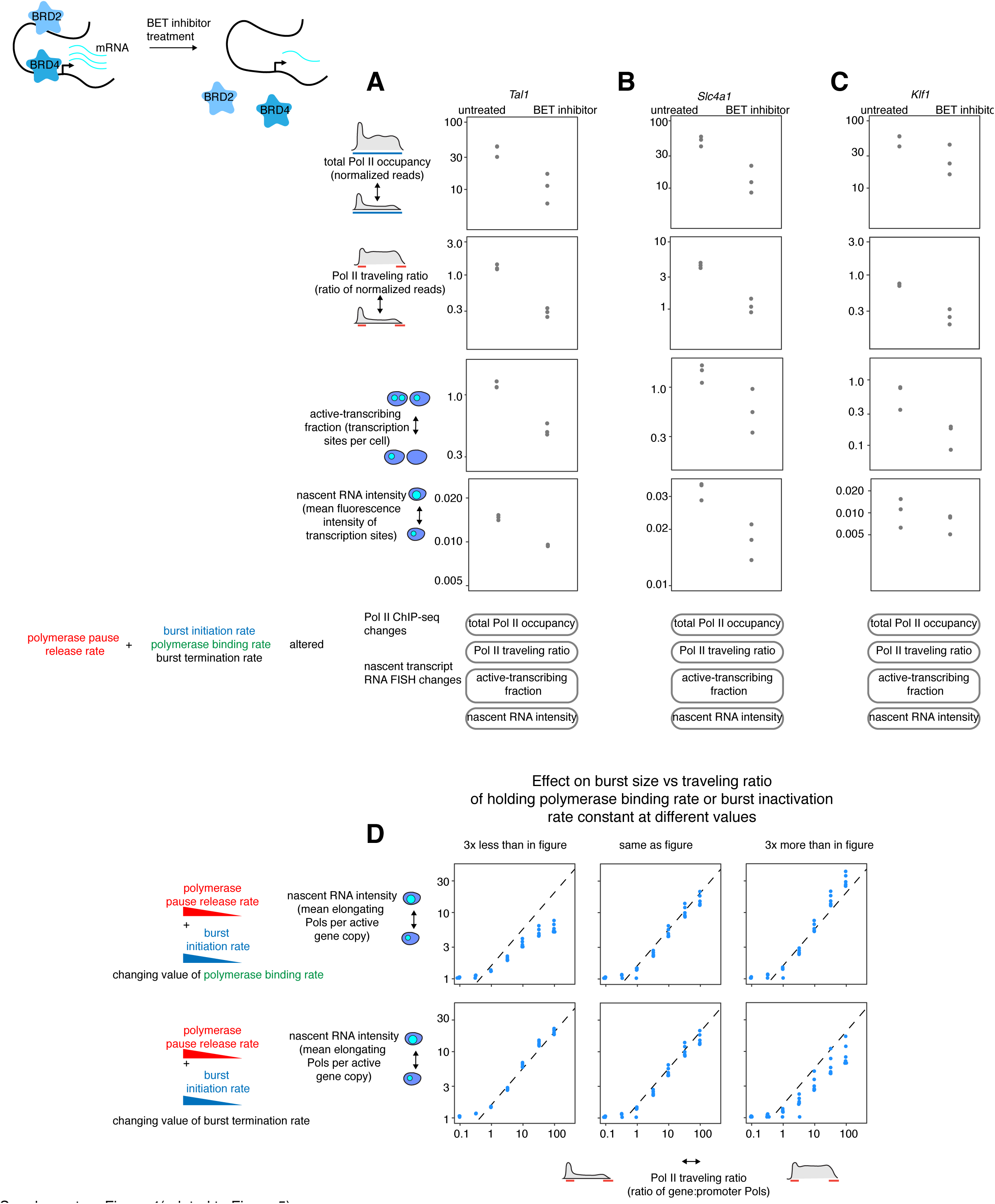
A. Pol II ChIP-seq and nascent transcript RNA FISH measurement of Tal1, untreated or with 60min 250nM JQ1. B. Pol II ChIP-seq and nascent transcript RNA FISH measurement of Slc4a1, untreated or with 60min 250nM JQ1. C. Pol II ChIP-seq and nascent transcript RNA FISH measurement of Klf1, untreated or with 60min 250nM JQ1. D. Effect of holding polymerase binding rate, or burst termination rate, constant at different levels on the relationship between nascent RNA intensity and traveling ratio when polymerase escape rate and burst initiation rate are changed.

**Supplementary Figure 5 (related to Figure 6):**
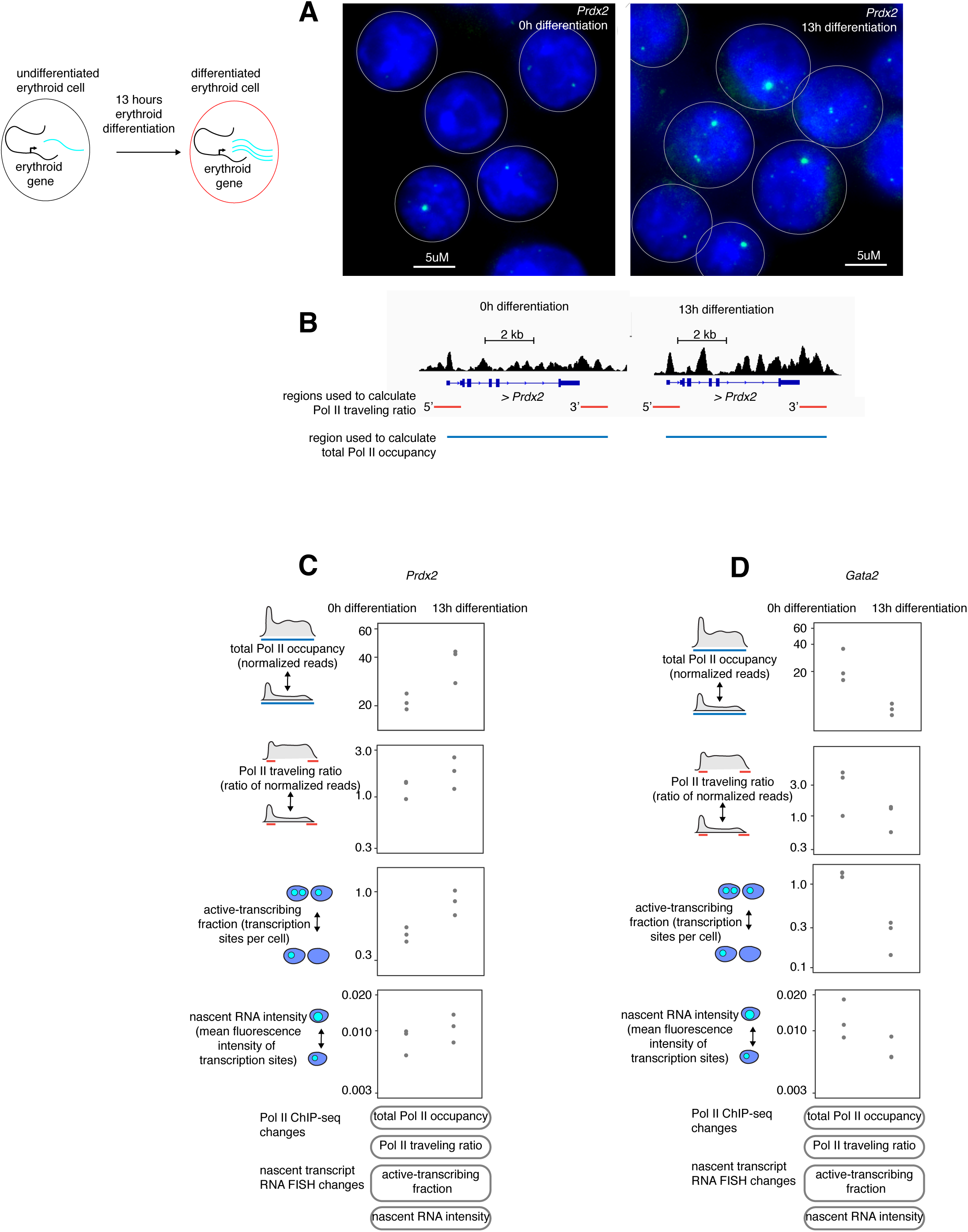
A. Intron RNA FISH images of Prdx2, undifferentiated or with 13h differentiation. B. Pol II ChIP-seq track of Prdx2, undifferentiated or with 13h differentiation (track heights normalized by library size). C. Quantification of total Pol II occupancy and Pol II traveling ratio and active-transcribing fraction and nascent RNA intensity for Prdx2, undifferentiated or with 13h differentiation, n=3 biological replicates. D. Quantification of total Pol II occupancy and Pol II traveling ratio and active-transcribing fraction and nascent RNA intensity for Gata2, undifferentiated or with 13h differentiation, n=3 biological replicates.

**Supplementary Figure 6(related to Figure 7):**
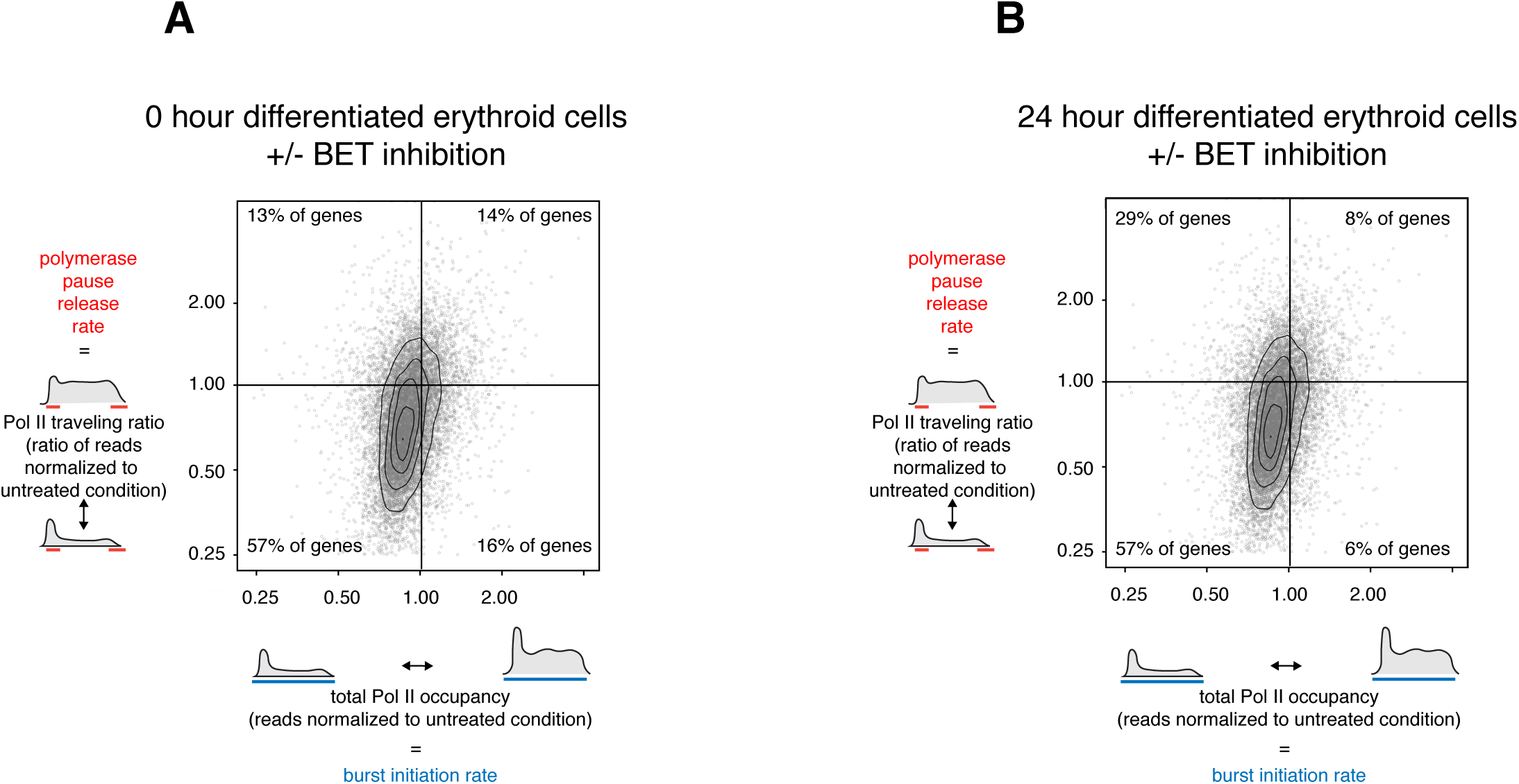
A. Pol II traveling ratio vs. total Pol II occupancy for all expressed genes in undifferentiated G1E-ER4 cells, untreated or with 60min 250nM JQ1, normalized to untreated cells. n=3 biological replicates. B. Pol II traveling ratio vs. total Pol II occupancy for all expressed genes in 24 hour differentiated G1E-ER4 cells, untreated or with 60min 250nM JQ1, normalized to untreated cells. n=3 biological replicates.

